# Epigenetic Control of Temperature-Dependent Female Reproductive Life History Trade-Offs in Seed Beetles, *Callosobruchus maculatus*

**DOI:** 10.1101/2021.10.08.463711

**Authors:** Beth A McCaw, Aoife M Leonard, Tyler J Stevenson, Lesley T Lancaster

## Abstract

Many species are threatened by climate change and must rapidly respond to survive changing environments. Epigenetic modifications, such as DNA methylation, can facilitate plastic responses by regulating gene expression in response to environmental cues. Understanding epigenetic responses is therefore essential for predicting species’ ability to rapidly adapt in the context of global environmental change. Here, we investigated the functional significance of DNA methylation on temperature-dependent life history in seed beetles, *Callosobruchus maculatus*. We assessed changes in DNA methyltransferase (*Dnmt1* and *Dnmt2*) expression levels under ambient conditions and thermal stress, and reproductive performance following artificially-induced epimutation via 3-aminobenzamide (3AB) and Zebularine (Zeb), at a range of ambient and warmer temperatures over two generations. We found that *Dnmt1* and *Dnmt2* were greatly expressed in females, throughout the body, and exhibited temperature-dependence; in contrast, *Dnmt* expression was minimal in males. Epimutation led to shifts in female reproductive life history trade-off allocation, and differentially altered thermal optima of fecundity and offspring viability. This study revealed the optimal allocation strategy among these fitness components is temperature-dependent, and trade-offs become increasingly difficult to resolve epigenetically under more extreme warming. Results suggest that epigenetic mechanisms are strongly implicated in, and perhaps limiting of, invertebrate life history responses to temperature change. Further investigation will reveal targeted DNA methylation patterns and specific loci associated with temperature-dependent life history trade-offs in seed beetles and other invertebrates.

## INTRODUCTION

Anthropogenic activities are causing rapid global climatic changes at unprecedented rates [1]. Many species must rapidly adjust their phenotype in order to survive the changing environment. While evolved responses are important, many adaptive responses may be additionally mediated through physiological or behavioural plasticity in life history (such as individual shifts in reproductive allocation strategies or dispersal abilities), thermal acclimation, or a combination of both [2]. Important life history traits such as fecundity and offspring viability often exhibit plasticity in response to temperature, such that temperature stress can mediate plastic shifts in life history allocation that are often adaptive [3, 4]. However, due to underlying trade-offs among different life history traits [5], a beneficial change in one such trait may cause a detrimental change in another trait, such that environmentally-dependent life history shifts, even while adaptive in one life history dimension, may have complex consequences for multiple traits and overall fitness [6]. Therefore, optimal life history allocation functions may themselves be temperature-dependent, although this possibility has received little attention to date.

Adjustment of life history allocation is known to be an important climate change response. This can occur when an environmental change induces shifts in energy allocation, such as increasing the need for resource acquisition due to temperature-dependent metabolic rates, or increased stress resistance due to thermal acclimation, which often leads to cascading shifts in growth or survival [7–12]. Life history allocations may also be adaptively regulated in response to disproportional changes in temperature-dependent fitness at different life stages. For instance, decreased offspring survival under thermal stress may lead to adaptive adjustments in fecundity or timing of reproduction in adulthood [13]. Such plastic, short- or long-term, environmentally induced trait shifts may increase the organism’s potential to adapt to or buffer against environmental change [14–17] or may be maladaptive, depending on population demography, the magnitude of environmental change and the resulting selection pressure and intensity of conflict in resource allocation [6, 18, 19]. Some studies have discovered physiological shifts affecting life history trade-offs under climate change [9, 11, 12, 13], highlighting the importance of deepening our understanding of the underlying mechanisms behind life history trade-offs, particularly those that are environmentally sensitive, to determine whether trade-offs will facilitate (compensatory mechanisms) or constrain (hindering mechanisms) species’ adaptive responses to climate.

Epigenetic mechanisms have previously been implicated in plastic responses to environmental change, including temperature change (reviewed in [20]), and have also been implicated in several critical life history allocation decisions [21–23]. Epigenetic mechanisms thus represent important candidates for adaptively matching life history trade-off structures to temperature. Epigenetic mechanisms in general depend on a set of molecules that modify DNA accessibility to transcription factors, and thus regulate gene transcription without directly altering the DNA sequence [24]. One such epigenetic mechanism, DNA methylation, is well known to regulate gene expression, although functionality can vary across taxa [25, 26]. In the best described model for DNA methylation functionality, derived primarily from vertebrates, the process of DNA methylation begins with the DNA (cytosine-C5) methyltransferase 3 enzyme, DNMT3 (encoded by the *Dnmt3* gene) – which *de novo* methylates the genomic template [27]. DNMT1 (encoded by *Dnmt1*) maintains DNA methylation on the genome [28, 29], while DNMT2 (encoded by *Dnmt2*) methylates tRNA [30].

The role of DNA methylation to regulate phenotypic responses to various environmental stressors has been widely studied across taxa, including in invertebrates [20, 31–33], and evidence suggests that this mechanism is well suited to facilitating adaptive plasticity of ecologically important traits, facilitating transmission of these phenotypes to future generations, and affecting population fitness and local adaptation. Environmental control of DNA methylation is likely to be particularly important in ectotherms, such as insects, whose internal temperatures are highly linked to changing external temperatures, and must therefore rapidly adjust thermally-dependent performance and life history strategies as their body temperature shifts [34].

The presence and function of DNA methylation and its associated enzymes are highly varied among insect taxa, and the relationship between DNA methylation and phenotypic plasticity in insects remains an active area of research [35–38]. Temperature induced changes in *Dnmt* expression or DNA methylation levels have previously been found in many insect species ([20] and references therein), and previous studies have further identified a role for DNA methylation in insect life history and fitness [21–23]. Here, in an insect model species, we evaluate temperature dependence of DNA methylation, and we further manipulate epigenetic potential to assess downstream consequences for temperature-mediated plasticity in life history traits. These combined approaches reveal complex, thermally-dependent epigenetic mediation of ectotherm life history traits and trade-off allocation strategies.

### Study System

The bruchid seed beetle, *Callosobruchus maculatus*, is a global crop pest species that uses a variety of leguminous crops as their obligate larval hosts. Its rapid life cycle and robust nature provides an effective, easily manageable insect model system for investigating the role of temperature to regulate DNA methylation, and the role of DNA methylation to mediate thermally-dependent expression of life history fitness components. Moreover, its ability to rapidly colonise a wide variety of global climate regions implies a high level of adaptive thermal performance plasticity and evolvability in response to novel thermal regimes [39], which has also been recently experimentally demonstrated [40]. Additionally, preliminary studies assessing *C. maculatus* thermal performance of fitness components under a range of rearing temperatures found that fecundity, offspring viability and generation time each displayed different thermal optima ranging from 25-32°C (Leonard & Lancaster, Unpublished Data), suggesting that optimal trade-off strategies among these traits may be temperature-dependent. Similarly, Berger *et al*. [41] found a temperature-dependent trade-off between longevity and reproductive effort in thermally stressed *C. maculatus* associated with improved germline repair of stress-induced DNA mutations – which may have strong implications on adaptive abilities to climate change. These earlier results suggest that temperature-dependent, plastic shifts in life history trade-offs may facilitate adaptive responses to climate warming in our study organism.

To test our prediction that DNA methylation influences temperature-dependent life history trait expression and optimal life history trade-off allocations, first we examined sex and anatomical differences in *Dnmt* gene expression to test for sex-specific differences in expression of enzymes that initiate DNA methylation, and also whether these represent whole organism differences or anatomical region specificity. We then examined temperature-dependent *Dnmt* expression, and assessed shifts in four life history parameters in *C. maculatus* under pharmaceutically altered DNA methylation at a range of ambient to heat stressful conditions. Together, this set of experiments suggest that temperature-dependent epigenetic modifications and downstream phenotypic consequences may allow insects to respond and adapt to a changing environment in the global context of climate change, at least under moderate temperature increases.

## MATERIALS AND METHODS

### Stock Culture

*Callosobruchus maculatus* are an invasive pest that have evolved to complete its entire developmental cycle within stored, dried leguminous agricultural products [39]. Female *C. maculatus* lay their eggs on the legume surface, the eggs hatch and the larvae burrow into the bean where they feed and complete their life cycle. The adults then emerge for reproduction. It takes approximately three weeks to complete the life cycle, depending on the external (thermal) environment [42]. Our stock culture of *C. maculatus* was provided from a population obtained from Niger and sustained in a large, outbred population by Paul Eady at the University of Lincoln for over 300 generations on cowpeas (*Vigna unguiculata*, ancestral host) at 27°C with a 12L:12D photoperiod (P. Eady, pers. comm) before being transferred to the University of Aberdeen and reared in large subpopulations under the same conditions since 2015. The stock culture and F_0_ generation of *Callosobruchus maculatus* were similarly reared at 27°C in a programmable incubator (LMS model 280NP refrigerated incubator; Kent, UK) on dried cowpeas with a 12L:12D photoperiod.

### Study 1 – Sex, Anatomical Localisation and Temperature Effects of *Dnmt* Gene Expression

#### Sex and body plan

To examine potential sex and anatomical differences in *Dnmt* expression, *Dnmt1* and *Dnmt2* expression was compared in untreated adult male and female *C. maculatus*. The putative *de novo* methylation gene, *Dnmt3*, was not studied as *C. maculatus* does not appear to express *Dnmt3* [43]. Although *Dnmt2* is not directly involved in DNA methylation, we were nevertheless interested in examining sex and temperature effects on tRNA methylation, to inform future studies of its role in insects. Genes that are not epigenetic enzyme encoding genes were also examined, to create an overall picture of how epigenetic genes may be responding similarly or differently to other genes with divergent functions in thermal stress (heat shock proteins 70 (*Hsp70*) and 90 (*Hsp90*); [44]) and life history (vitellogenin (*Vtg*); [45]). Three replicates of six pooled, whole (non-dissected) individuals of each sex were obtained from the stock culture for RNA extraction. We also collected three replicates of six pooled heads and three replicates of six pooled bodies for each sex, to test for localised patterns within each sex. The comparison of head vs. body was intended as an initial approach to identify whether regional specificity in *Dnmt* expression might occur, for instance reflecting overrepresentation in the brain or reproductive tissues. Beetles were immediately frozen at -70°C until gene expression was evaluated by quantitative PCR (qPCR, see Methods below).

#### Temperature-dependent Dnmt expression

The results of our sex-body plan study suggested that *Dnmt1* and *Dnmt2* were primarily expressed throughout adult females (see Results). This finding, and our goal to understand the role of DNA methylation specifically for female reproductive trade-offs, led us to limit our temperature-dependent assay (and whole-genome methylation study) to whole adult females. To determine the effects of temperature on *Dnmt* expression in *C. maculatus*, six replicates of four pooled adult females were collected from the temperatures 22°C, 27°C, 31°C and 35°C. Beetles were cultured in thermally stable water baths at 31°C and 35°C with a natural daylight, while beetles cultured at 22°C and 27°C were kept in set incubators with a 12L:12D artificial light. Samples were stored at -70°C until RNA extraction and characterisation of *Dnmt1* and *Dnmt2* expression via qPCR.

#### Quantification of RNA expression

Pooled samples were homogenised mechanically, and RNA was extracted using a standard Trizol-chloroform procedure (ThermoFisher Scientific; [46]). RNA concentration and quality were determined using Nanodrop spectrophotometry (ThermoFisher Scientific). RNA samples were then washed with RNase-free DNase water (Invitrogen, ThermoFisher Scientific). cDNA was synthesized using 2.0µg of each RNA sample in a 20.0µL reaction with an oligo(dT)_18_ primer, according to the instructions provided by nanoScript2 Reverse Transcription Kit (Primerdesign Ltd). cDNA was stored at -20°C until its use in qPCR [46].

During the qPCR preparation stage, iQ Sybr Green Supermix (Bio-Rad Laboratories) was used to quantify targeted RNA expression levels via amplification. Each Supermix contained: 500µl Sybr Green, 200µl RNase-free DNase water and 50µl of a chosen primer. 15µl of Supermix was used for each reaction with 5µl of cDNA for a total volume of 20µl. All qPCR runs were performed in technical duplicate. Primers for target and positive control genes were designed by Primerdesign Ltd and Invitrogen, ThermoFisher Scientific, based on sequence data identified from the *C. maculatus* annotated genome ([47]; S4 Table). *18S rRNA* was used as a reference for all target and control gene expression calculations and was amplified using a previously published primer sequence ([48]; S4 Table). Quantitative PCRs were performed using a Bio-Rad Laboratories CFX96 system and were evaluated using a quantification stage and a melting curve specificity stage. The quantification stage involved denaturing cDNA at 95°C for 30 seconds, followed by 39 cycles of 1) 95°C for 30 seconds, 2) annealing of primers at their annealing temperatures for 30 seconds (S4 Table) and 3) cDNA extension at 72°C for 30 seconds. The number of cycles until amplification, detected by Sybr Green fluorescence, determined the cycle thresholds (Ct). At the melting curve specificity stage, the temperature decreased to 55°C for 5 seconds, then gradually increased by 0.5°C every second until it reached 95°C. The temperature at which the cDNA denatures verifies whether qPCR amplified one specific product. Real-time PCR Miner Software [49] was used to calculate the PCR efficiencies and Ct values from the raw fluorescence data. The fold expressions of *Dnmt*s, *Hsp*s and *Vtg* were measured in relation to the average cycling times of *18S rRNA* reference gene using 2^−ΔΔCt^ [50].

### Study 2 – Pharmaceutical Manipulation of DNA Methylation on Life History Components

Epigenetic variation was induced in treated beetles through artificial epimutations involving two DNA methylation manipulation drugs, each with opposing effects on DNA methylation *in vivo*. 3-aminobenzamide (3AB) (Sigma-Aldrich) induces DNA hypermethylation by inhibiting poly(ADPribose) polymerase and consequently inhibiting poly(ADP-ribosyl)ation, an important mechanism that prevents full methylation of the genome, thus allowing DNMT1 to methylate previously unmethylable cytosines post-replication [51, 52]. Conversely, Zebularine (Zeb) (Selleckchem, Houston, TX, USA) acts as a DNA methylation inhibitor. It has a high affinity for DNA methyltransferases and forms a covalent bond with carbon 6 of the DNMT. This inhibits the reaction pathway of DNMT catalysing the transfer of the methyl group from S-adenyl methionine to cytosine-C5 [53]. Preliminary studies confirmed that 125nM Zeb resulted in maximal effects on *C. maculatus* reproductive performance (where 12.5nM, 125nM, 500nM and 5µM concentrations were assessed; Moore, Stevenson & Lancaster, Unpublished Data). An additional pilot study confirmed that 125nM concentration of 3AB and Zeb were both maximally effective to induce changes in thermal performance in a different insect species (Areshi & Lancaster, Unpublished Data), so we prepared and used this concentration for both drugs, administered in the same way as the *C. maculatus* preliminary study. Drugs were initially dissolved in DMSO and then diluted in deionised water. Cowpeas were soaked in each treatment to ensure the delivery of treatment into the beetles through ingestion during the larval stage. After soaking in Zeb, 3AB or deionised water for 24 hours, the beans were dried in an oven at 60°C to return moisture levels similar to that of normal dried beans.

To determine whether DNA methylation influences life history trait allocation in response to warming, *C. maculatus* beetles were exposed to one of three treatments: deionised water (control treatment), 3AB or Zeb, and were cultured at 27°C, 31°C or 35°C for two generations. Each replicate consisted of a female beetle paired with a random male from the F_0_ stock culture and placed in a vial or petri dish with 10g of beans on which to oviposit. Upon adult emergence of F_1_ generation, at least two virgin males and two virgin females were taken from each replicate to generate 13 F_1_ mated pairs per treatment x temperature combination. Pairing was carried out in each treatment x temperature combination by mating females with males from different replicate vials to prevent pairing of siblings. These pairs were each established on 10g of fresh beans of the same treatment and cultured in the same temperature as their parents had experienced. Data collection on beetle performance included F_1_ fecundity (total number of eggs), F_2_ viability (total number of F_2_ emerged adults divided by total number of eggs laid by F_1_), F_2_ developmental period (number of days from egg to adult, averaged within F_1_ families) and F_2_ adult lifespan (also averaged within F_1_ families). We also investigated temperature and epimutation treatment effects on whole-genome methylation using methylation-sensitive ELISAs (Study in Supporting Information).

### Statistical Analysis

#### Gene expression

All statistical analyses were performed in R version 3.6.3 and figures were plotted using the “ggplot2” package [54, 55]. Shapiro-Wilk normality tests were used to assess data normality. First, after log transformation of *Dnmt1* and *Dnmt2* data established normality, linear models examined sex differences in expression of *Dnmt1* and *Dnmt2* in untreated, non-thermally challenged (ancestral 27°C) whole beetles. Second, nested linear models were used to examine whether *Dnmt1* and *Dnmt2* expression varied according to body plan (head vs. body) within each sex. Linear models and nested linear models were also used to examine sex and body plan differences in expression of *Hsp70, Hsp90* and *Vtg*, with *Vtg* expression data log transformed to meet linear model assumptions. Third, linear models were used to examine the effects of temperature on *Dnmt1* and *Dnmt2* expression in adult females (*Dnmt2* data log transformed), with temperature fitted as an orthogonal polynomial term (with the poly() function in R) which independently assesses first order and quadratic effects of temperature after accounting for autocorrelation among main and quadratic terms.

#### Life history traits

Life history performance variables were each modelled as a function of drug treatment nested within each temperature, to reveal the experimental temperatures at which epimutation has a significant effect. F_1_ fecundity was modelled fitting a general linear model with a quasipoisson error distribution, as this represented count data with a large dispersion parameter (6.444553). F_2_ offspring viability was fitted using a quasibinomial error distribution (dispersion parameter = 5.947479). F_2_ developmental period and adult lifespan both conformed to assumptions of normality and thus linear models were applied. Effects of drug were assessed with water-treated controls as the baseline within each temperature regime, using nested (general) linear models.

## RESULTS

### Study 1 – Sex, Anatomical Localisation and Temperature Effects of *Dnmt* Gene Expression

#### Dnmt1

Expression (fold change with respect to the *18S rRNA* reference gene) was significantly greater in whole (non-dissected) adult females than in adult males (effect of female sex = 1.66 ± 0.57 SE, t = 2.89, P = 0.04; *n* = 6; Figure 1A). Sex differences in *Dnmt1* expression were reproducibly detected when comparing expression in the different body parts (main effect of sex in body part assay = -2.60 ± 0.93 SE, t = -2.78, P = 0.03; *n* = 11), however there was no significant difference in *Dnmt1* expression between the body parts for either males (effect of head vs body, nested in males = - 0.19 ± 0.83 SE, t = -0.22, P = 0.83) or females (effect of head vs body, nested in females = -0.99 ± 0.93 SE, t = -1.06, P = 0.32), indicating that elevated *Dnmt1* detected in adult females (in comparison to males) is expressed throughout the female body and not only in, for example, her brain or developing eggs. In addition, there was a significant positive linear relationship of *Dnmt1* expression in whole adult females in response to temperature (effect of poly(temp,2)1: 2.47 ± 0.57 SE, t = 4.35, P = 0.00028; effect of poly(temp,2)2: -0.55 ± 0.57 SE, t = -0.97, P = 0.35; *n* = 24; Figure 2A).

**Fig. 1.**
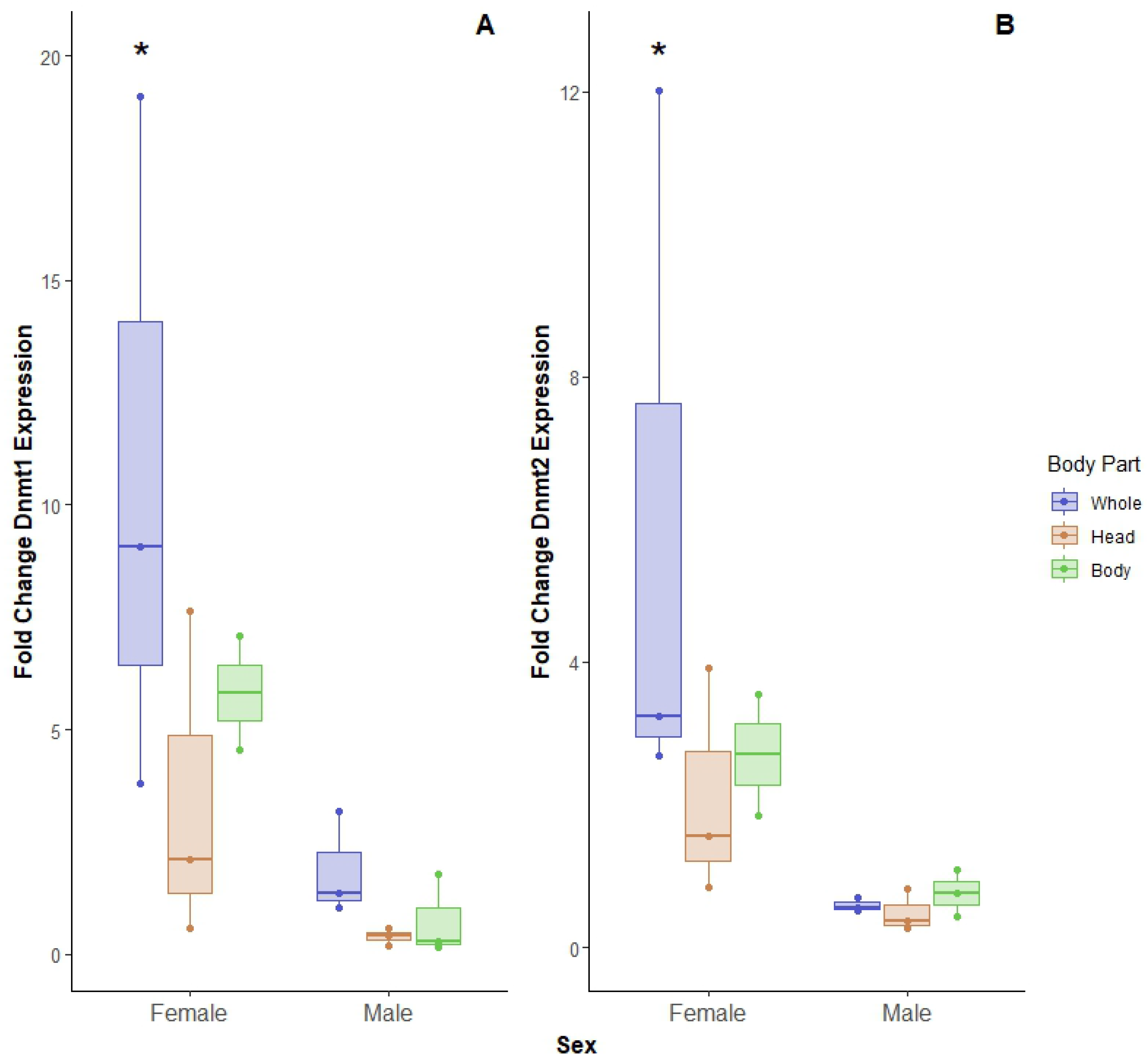
**Fold change in RNA expression of A) *Dnmt1* and B) *Dnmt2* in adult female and adult male whole body (blue, left; *n* = 6) and head (orange, middle) vs body (green, right) (*n* = 11), normalised over the expression of *18S rRNA* reference gene and determined by qPCR.** Data was back transformed for visualisation. Asterisks indicate significant differences in expression in whole females compared to whole males. Each box represents the median, lower 25^th^ and upper 75^th^ percentile, with whiskers showing the values within 1.5 times the interquartile ranges. Outside values are >1.5 times and <3 times beyond the interquartile ranges.

**Fig. 2.**
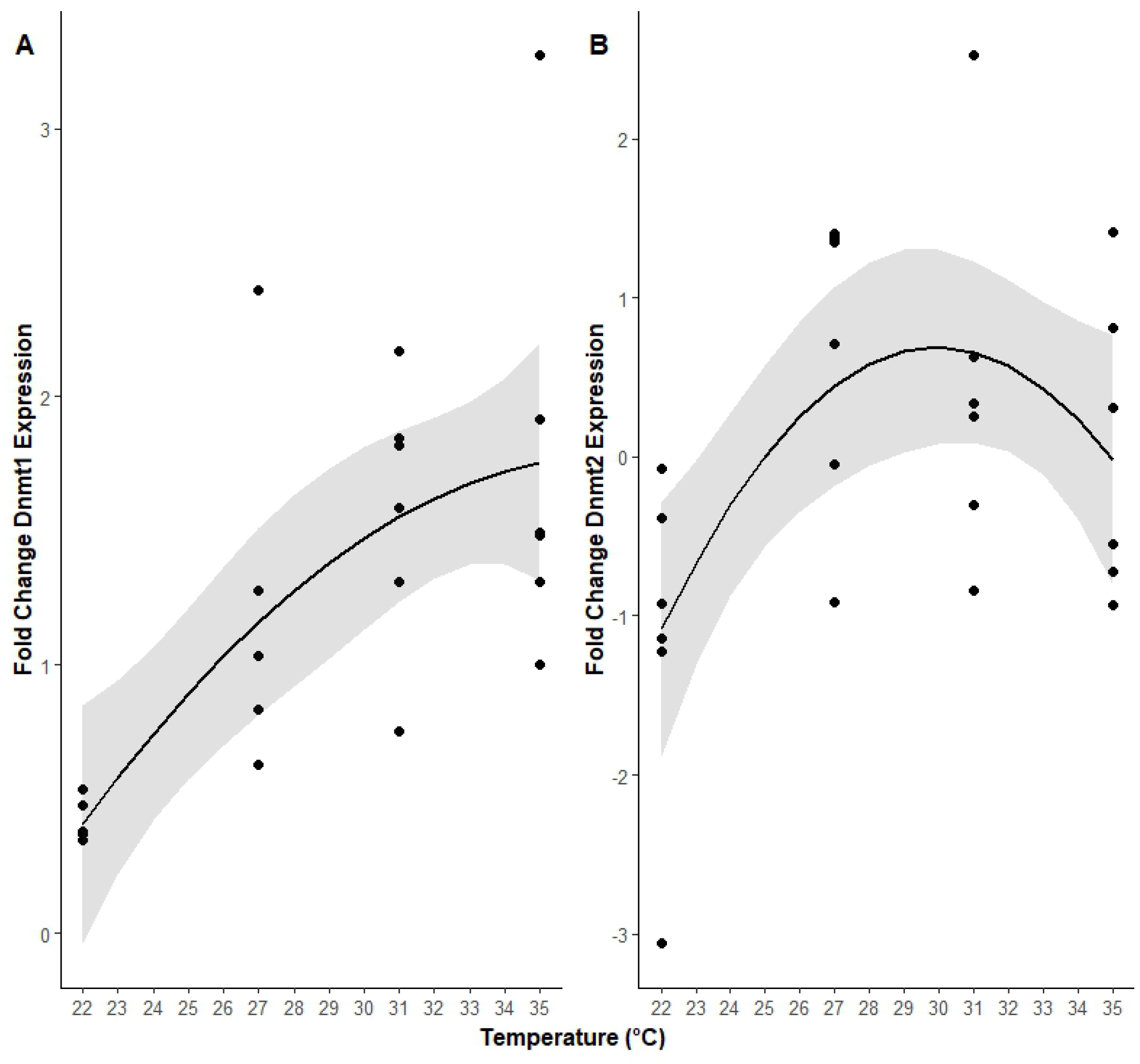
**Fold change in A) *Dnmt1* and B) *Dnmt2* expression levels in adult female life stage across temperatures, normalised over the expression of *18S rRNA* reference gene and determined by qPCR (*n* = 24).** Shaded regions represent 95% confidence intervals.

#### Dnmt2

Fold change expression was significantly greater in whole adult females than in adult males (effect of female sex = 2.08 ± 0.48 SE, t = 4.34, P = 0.01; *n* = 6; Figure 1B). Sex differences in *Dnmt2* expression were reproducibly detected when comparing expression in body parts (main effect of sex in body part assay: -1.29 ± 0.54 SE, t = - 2.38, P = 0.05; *n* = 11), however *Dnmt2* expression was not significantly different between head and body of adult females (effect of head vs body, nested in females = -0.39 ± 0.54 SE, t = -0.73, P = 0.49) or males (effect of head vs body, nested in males = -0.49 ± 0.48 SE, t = -1.01, P = 0.35). Linear models revealed a significant quadratic change in *Dnmt2* expression in whole adult females across temperatures (effect of poly(temp,2)1: 2.00 ± 1.01 SE, t = 1.97, P = 0.06; effect of poly(temp,2)2: -2.61 ± 1.01 SE, t = -2.57, P = 0.02; *n* = 24; Figure 2B), suggesting that *Dnmt2* expression is also temperature-dependent in females, but differently so to *Dnmt1*.

#### Other genes

Linear models and nested linear models revealed no significant sex or body plan differences in fold change expression of *Hsp70, Hsp90* and *Vtg* with respect to the reference gene (S5 and S6 Tables), providing additional confirmation that sex differences in expression of *Dnmt1* and *Dnmt2* are specific to epigenetic machinery, and not merely reflective of sex differences in expression of (other) environmentally or developmentally responsive genes.

### Study 2 – Pharmaceutical Manipulation of DNA Methylation on Life History Components

#### Fecundity

Nested linear models demonstrate an overall main effect of temperature on fecundity (main effect of 31°C in comparison to 27°C: -0.30 ± 0.15 SE, t = -1.97, P = 0.05; main effect of 35°C: -0.73 ± 0.17 SE, t = -4.25, P = 4.51e-05). Overall, fecundity declined as a function of temperature, but drug treatment affected the form of this relationship, as Zeb treatment increased fecundity at 35°C (Table 1; Figure 3A). Neither methylation drug treatment had an effect on fecundity at 31°C in comparison to the control treatment. Separate analysis of drug treatment effects at 27°C only confirm that fecundity was significantly lower for 3AB treatment in comparison to the control treatment (effect of 3AB at 27°C only: -0.24 ± 0.09 SE, t = -2.70, P = 0.01), while Zeb treatment continued to have no effect on fecundity at 27°C (effect of Zeb: 0.03 ± 0.08 SE, t = 0.32, P = 0.75).

**Table 1:**
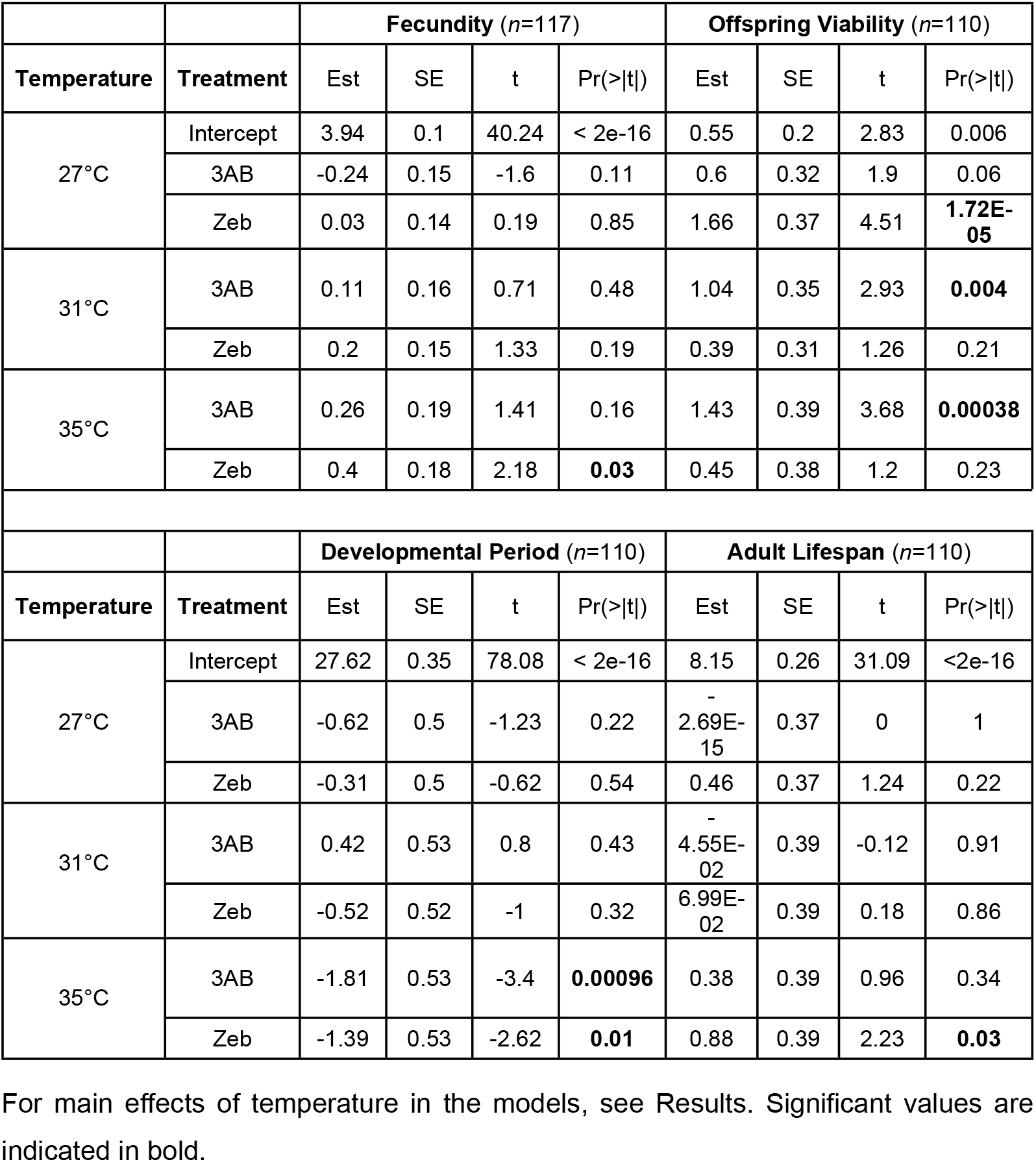
**Nested models explaining effects of treatment on fecundity, offspring viability, developmental period and adult lifespan at different temperatures (baseline = H2O)**.

|  |  | Fecundity (*n*=117) |  |  |  | Offspring Viability (*n*=110) |  |  |  |
| --- | --- | --- | --- | --- | --- | --- | --- | --- | --- |
| Temperature | Treatment | Est | SE | t | Pr(>\|t\|) | Est | SE | t | Pr(>\|t\|) |
| 27°C | Intercept | 3.94 | 0.1 | 40.24 | < 2e-16 | 0.55 | 0.2 | 2.83 | 0.006 |
|  | 3AB | -0.24 | 0.15 | -1.6 | 0.11 | 0.6 | 0.32 | 1.9 | 0.06 |
|  | Zeb | 0.03 | 0.14 | 0.19 | 0.85 | 1.66 | 0.37 | 4.51 | **1.72E-05** |
| 31°C | 3AB | 0.11 | 0.16 | 0.71 | 0.48 | 1.04 | 0.35 | 2.93 | **0.004** |
|  | Zeb | 0.2 | 0.15 | 1.33 | 0.19 | 0.39 | 0.31 | 1.26 | 0.21 |
| 35°C | 3AB | 0.26 | 0.19 | 1.41 | 0.16 | 1.43 | 0.39 | 3.68 | **0.00038** |
|  | Zeb | 0.4 | 0.18 | 2.18 | **0.03** | 0.45 | 0.38 | 1.2 | 0.23 |

|  |  | Developmental Period (*n*=110) |  |  |  | Adult Lifespan (*n*=110) |  |  |  |
| --- | --- | --- | --- | --- | --- | --- | --- | --- | --- |
| Temperature | Treatment | Est | SE | t | Pr(>\|t\|) | Est | SE | t | Pr(>\|t\|) |
| 27°C | Intercept | 27.62 | 0.35 | 78.08 | < 2e-16 | 8.15 | 0.26 | 31.09 | <2e-16 |
|  | 3AB | -0.62 | 0.5 | -1.23 | 0.22 | -<br>2.69E-15 | 0.37 | 0 | 1 |
|  | Zeb | -0.31 | 0.5 | -0.62 | 0.54 | 0.46 | 0.37 | 1.24 | 0.22 |
| 31°C | 3AB | 0.42 | 0.53 | 0.8 | 0.43 | -<br>4.55E-02 | 0.39 | -0.12 | 0.91 |
|  | Zeb | -0.52 | 0.52 | -1 | 0.32 | 6.99E-02 | 0.39 | 0.18 | 0.86 |
| 35°C | 3AB | -1.81 | 0.53 | -3.4 | **0.00096** | 0.38 | 0.39 | 0.96 | 0.34 |
|  | Zeb | -1.39 | 0.53 | -2.62 | **0.01** | 0.88 | 0.39 | 2.23 | **0.03** |

For main effects of temperature in the models, see Results. Significant values are indicated in bold.

**Fig. 3.**
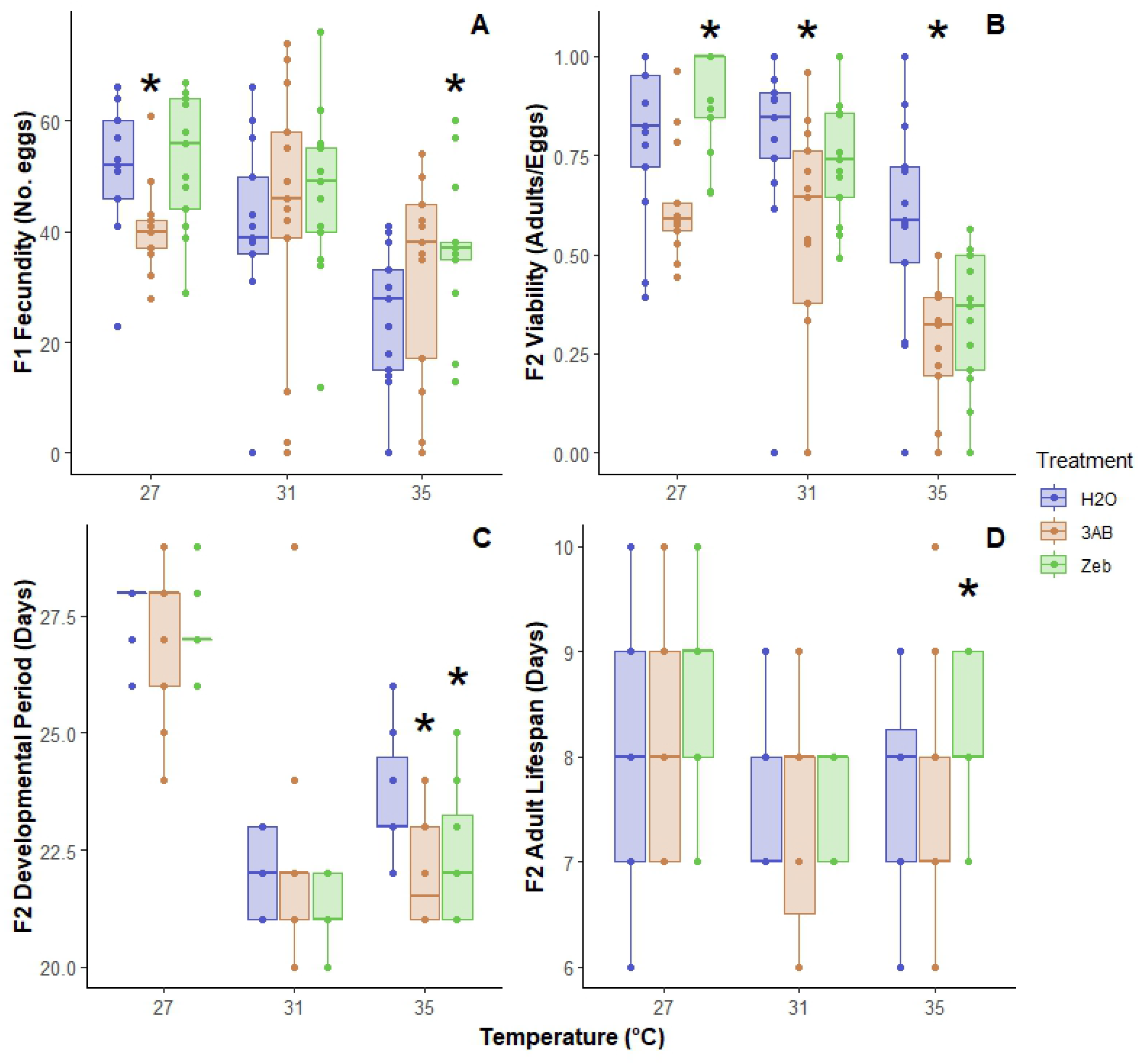
**Effects of temperature and DNA methylation drug treatment on A) fecundity (*n* = 117), B) offspring viability (*n* = 110), C) developmental period (*n* = 110) and D) adult lifespan (*n* = 110) at 27°C, 31°C and 35°C.** Within each temperature: water = blue, left; 3AB = orange, middle; Zeb = green, right. Asterisks indicate significant effects of drug treatment within each temperature category. Each box represents the median, lower 25th and upper 75th percentile, with whiskers showing the values within 1.5 times the interquartile ranges. Outside values are >1.5 times and <3 times beyond the interquartile ranges.

#### Offspring Viability

Elevated temperatures, particularly 35°C, had a significant effect on reducing offspring viability (in the nested model, main effect of 31°C in comparison to 27°C: -0.03 ± 0.30 SE, t = -0.11, P = 0.91; main effect of 35°C: -1.50 ± 0.36 SE, t = - 4.18, P = 0.000063), and drug treatment altered the form of this relationship as follows: Zeb treatment significantly increased offspring viability in comparison with the control treatment at 27°C, while 3AB had no significant effect. In contrast, 3AB treatment significantly increased offspring viability at elevated temperatures (31°C and 35°C), with no effect of Zeb at these temperatures (Table 1; Figure 3B).

#### Developmental Period

Temperature had a significant main effect of reducing developmental period (in the nested model, main effect of 31°C in comparison to 27°C: -5.71 ± 0.52 SE, t = -10.92, P = < 2e-16; main effect of 35°C: -3.89 ± 0.52 SE, t = - 7.44, P = 3.38e-11). No significant effect of drug treatment was detected on developmental period at 27°C and 31°C in comparison with the control treatment. However, developmental period was significantly decreased by both 3AB and Zeb at 35°C (Table 1; Figure 3C).

#### Adult Lifespan

Temperature did not significantly affect adult lifespan (in the nested model, main effect of 31°C in comparison to 27°C: -0.61 ± 0.39 SE, t = -1.57, P = 0.12; main effect of 35°C: -0.70 ± 0.39 SE, t = -1.81, P = 0.07). There was no significant difference in adult lifespan for either Zeb or 3AB treatment in comparison to water treatment at 27°C and 31°C. Zeb treatment increased adult lifespan at 35°C, although effects are very subtle (Table 1; Figure 3D).

## DISCUSSION

Understanding the underlying mechanisms of life history allocation shifts in response to temperature change is essential for understanding the ability of organisms to rapidly adapt to novel climates. Evidence of temperature-dependent *Dnmt* expression in adult females and evidence of the (divergent) influence of experimentally altered patterns of DNA methylation on fecundity, offspring viability, developmental period, and adult lifespan all support the hypothesis that DNA methylation is involved in regulating shifts in female life history strategies and fitness at different temperatures.

### Dnmt Expression

*Dnmt1* and *Dnmt2* expression levels were greater in adult females than in males (Figure 1), suggesting that *Dnmt*s have sex-specific roles, and may particularly help regulate female functions, for instance by regulating DNA methylation levels to match life history patterns to thermal conditions (see below). Previous studies have also investigated the role of DNA methylation in female reproductive life history strategies. For instance, a recent study found that gene knockdown of *Dnmt1* and methyl-DNA-binding protein 2/3 (*MBD2/3*) - a gene involved in gene body methylation in insects - and methylation inhibition by 5-aza-2’-deoxycytidine (a pharmaceutical alternative to Zeb) impaired ovary development and reduced fecundity and hatch rates in developing silkworm (*Bombyx mori*), tobacco cutworm (*Spodoptera litura*) and migratory locust (*Locusta migratoria*), suggesting intragenic DNA methylation regulates insect reproduction [56]. Silencing the Dnmt1 associated protein 1 (*DMAP1*) gene severely reduced fecundity and offspring survival in the harlequin ladybird (*Harmonia axyridis*), suggesting a role of the *DNMT-DMAP1* complex in life history traits in *H. axyridis* [23]. Similarly, *Dnmt* silencing reduced the number of offspring produced in the brown plant hopper (*Nilaparvata lugens*) [21] and reduced egg production and embryo viability in the large milkweed bug (*Oncopeltus fasciatus*) [22]. Together, these results demonstrate an evolutionary conserved role of DNA methylation in female reproductive life history strategies across insects.

To our knowledge, present results are the first evidence of temperature-dependent *Dnmt1* and *Dnmt2* expression in an insect adult female life stage (Figure 2) and suggest that temperature dependency in *Dnmt* expression functions in the epigenetic regulation of temperature-dependent reproductive life history strategies in insects (see below). Our finding of temperature dependence in the expression of the tRNA methylation enzyme-encoding gene, *Dnmt2*, is particularly interesting given the limited knowledge on *Dnmt2*, emphasizing the need for more research to better understand the role of tRNA methylation in response to temperature.

Our results indicate that there was no significant difference in *Dnmt* expression observed between the head and body of adult females. This suggests that expression is throughout the adult female and not just in her eggs and ovaries, providing evidence that DNA methylation is involved in other non-gamete development processes. The comparison of head vs. body was intended as a primary method to identify anatomical localisation of *Dnmt1* and *Dnmt2* expression, for instance reflecting overrepresentation in the brain or reproductive tissues. Having found no difference between heads and (headless) bodies, we are able to tentatively conclude that the differences observed among individuals reflect whole-organism shifts. These findings may contradict previous studies that suggest an additional role of DNA methylation enzymes in reproductive tissues and gametogenesis in insects. For instance, significantly greater *Dnmt1* expression was found in the oviposited eggs compared to the whole body of female adult red flour beetles (*Tribolium castaneum*), an insect that lacks a methylated genome but expresses *Dnmt1* [57] and in the ovaries compared to other tissues in developing *Bombyx mori* [56]. Similarly, in the milkweed bug, *Dnmt1* may play a specific role in oogenesis and is not required for somatic cell function (Amukamara *et al*., 2020). In addition, *Dnmt1* was highly expressed in the ovaries and in the testes of fire ants (*Solenopsis Invicta*) [58]. We did not measure *Dnmt* expression directly in the gonads, therefore, further studies on the *Dnmt* expression in all seed beetle life stages, including egg, all larval stages and pupae stages, and in reproductive tissues, as examined by Schulz *et al*. [57], Kay *et al*. [58] and Amukamara *et al*. [37], is required to better understand its role in reproductive development in insects. Collectively, this could provide insight into the functionality behind the variation in the amount of methylated genomic DNA and the expansion and losses of *Dnmt* subfamilies across insect taxa [35].

### Effects on Female Life History

Developmental period was significantly decreased by both 3AB and Zeb at 35°C (Figure 3C), suggesting that both treatments increased fitness at higher temperatures by reducing generation times, thereby widening the thermal optimum for fitness. Development time is generally optimal at 31°C in water-treated controls, and treatment appears to have induced this same (31°C) developmental time at 35°C, while control individuals suffered at 35°C with a greater generation time. This suggests that drug treatments may be influencing transcription of different loci associated with development at higher temperatures. Moreover, Zebularine treatment significantly increased adult lifespan at 35°C (Figure 3D), suggesting that Zebularine also increased thermal niche breadth via this life-history parameter at high temperatures. Overall, methylation drug treatments only influenced developmental period and adult lifespan at increasing temperatures, and not at the ancestral 27°C.

Fecundity overall declined at elevated temperatures, in comparison to the baseline 27°C. However, fecundity was improved at 35°C following Zebularine treatment compared to water treatment (Figure 3A), suggesting that the inhibition of DNA methylation increases thermal niche breadth via enhanced fecundity at increasing temperatures. In contrast, offspring viability was highest at 31°C in control females, and decreased at 35°C. However, an increase in offspring viability was observed following induced DNA hypermethylation by 3-aminobenzamide treatment at both 31°C and 35°C (Figure 3B). Increased offspring viability has been observed in many insects as an epigenetic response to temperature change [59–61], indicating epigenetic regulation of offspring viability may be important as a survival strategy for a changing climate, particularly in insects.

By comparing the effects of methylation drug treatment and temperature on fecundity and offspring viability, we find that, at ambient temperatures, artificial epimutation has opposing effects (where *hypo*methylation increased offspring viability, while *hyper*methylation decreased fecundity; this suggests lower levels of DNA methylation are overall favoured at ambient thermal conditions). In contrast, at elevated temperatures, the different epimutations always acted to increase fitness parameters, but each acted to increase a different life history parameter (*hypo*methylation increased the thermal breadth of fecundity, but *hyper*methylation increased thermal breadth of offspring viability). This latter result suggests that optimal DNA methylation levels for life history allocations are more difficult to achieve as temperature increases, and further suggests that life history trade-offs may be more intense and difficult to resolve as temperatures warm. We suggest that, in general, the performance of epigenetic perturbation across a range of temperatures, and monitoring of life history responses, may provide a revealing and helpful research tool to uncover life history trade-off functionality under predicted global warming.

It is well known that the genotype-environment interactions are a common feature of life history trade-offs [41], and that plastic shifts in trait allocation can therefore shape broader ecological strategies and syndromes to facilitate environmental adaptation [62]. By studying the epigenetic mechanisms and effects of epigenetic perturbations on this interaction, this study shows that DNA methylation is involved in regulating phenotypic plasticity in the form of resolving temperature-dependent fecundity-offspring viability fitness trade-offs in seed beetles under thermal change. Previous studies have also shown that DNA methylation is involved in fitness trade-offs in response to environmental stress [63–65]. However, present findings are the first to demonstrate that DNA methylation is involved in temperature-dependent female reproductive trade-offs. This provides further insight on how epigenetic mechanisms allow seed beetles, and possibly other insects of similar life-history strategies, to adapt to a changing climate.

Insects with different life-history strategies are predicted to respond differently to global warming. Multivoltine insects, like *C. maculatus*, with a rapid life cycle and no diapause stage, are likely to respond the greatest to increasing temperatures by increasing their distribution range, especially under increasingly benign conditions [66, 67]. With this prediction, present results suggest that environmental epigenetics in *C. maculatus* could increase their potential to expand their distribution range in response to increasing temperatures. This could occur through either expanding their geographical range or expanding their plant host species range. Many studies have shown that *Callosobruchus maculatus* can rapidly adapt to novel bean types. For instance, Price *et al*. [68] demonstrated that, within just five generations of adapting to novel hosts, the population fitness almost returns to its optimal state of fitness as when reared on the cowpea ancestral host. Collectively, to tolerate environmental stress, DNA methylation could provide individuals with opportunities to adjust different life history strategies which could explain their ability to rapidly adapt to novel host species and climates as they spread.

Our preliminary analyses of how whole-genome methylation is affected by temperature and treatment revealed suggestive trends but did not yield conclusive results likely due to low power to detect shifts in methylation against the backdrop of moderate sample sizes and high baseline individual variability in methylation levels (see power analysis in Supporting Information). Our analyses therefore suggest that changes in DNA methylation induced by temperature and treatment are subtle and locus-specific, despite being associated with profound phenotypic effects (see also Sarma *et al*. [69], where they show only a small change in genome-wide DNA methylation in response to Zebularine treatment but profound phenotype shifts). Further investigations are required using more fine-scale approaches to examine the DNA methylation pattern of targeted genes associated with phenotypic expression in seed beetles.

## CONCLUSION

The results of this study provide novel findings that DNA methylation plays a functional role in regulating *C. maculatus* female reproductive life history strategies and how they, and their optima, vary with temperature, reflected by offspring thermal performance. The results suggest that the observed temperature dependence of *Dnmt* expression in adult females, rather than being simply a thermodynamic property of the molecule, may reflect a highly evolved trade-off between different components of fitness at different temperatures, to which epigenetic mechanisms regulate temperature-dependent reproductive allocation decisions. Given that ectotherms have long evolved in the context of experiencing a wide range of body temperatures, each of which need to be optimised over evolutionary time in order to maximise fitness, it is likely that other ectotherms also have commonly evolved complex epigenetic regulatory mechanisms to evolve such temperature-dependent trade-offs. Further investigations will identify the location of DNA methylation and how these mechanisms regulate the expression of genes associated with the phenotypic fitness parameters in question. This line of research will provide a greater understanding of how insects respond and adapt to environmental change.

## ACKNOWLEDGEMENTS

Thank you to P. Eady for providing *C. maculatus* to initiate our laboratory population and advice on rearing them. Thank you to A. Sayadi and G. Arnqvist for their contributions to the *C. maculatus* transcripts. Thank you to Holly Marshall for commenting on an earlier version of the ms. Thank you to Elisabetta Tolla and Lana Dunan for their guidance and support throughout the study.

## SUPPORTING INFORMATION

**S1 Table. Number of replicate pooled samples (six adult females per pool) for each treatment x temperature combination for whole-genome methylation analysis, balanced over four ELISA plates**.

**S1 Fig.**
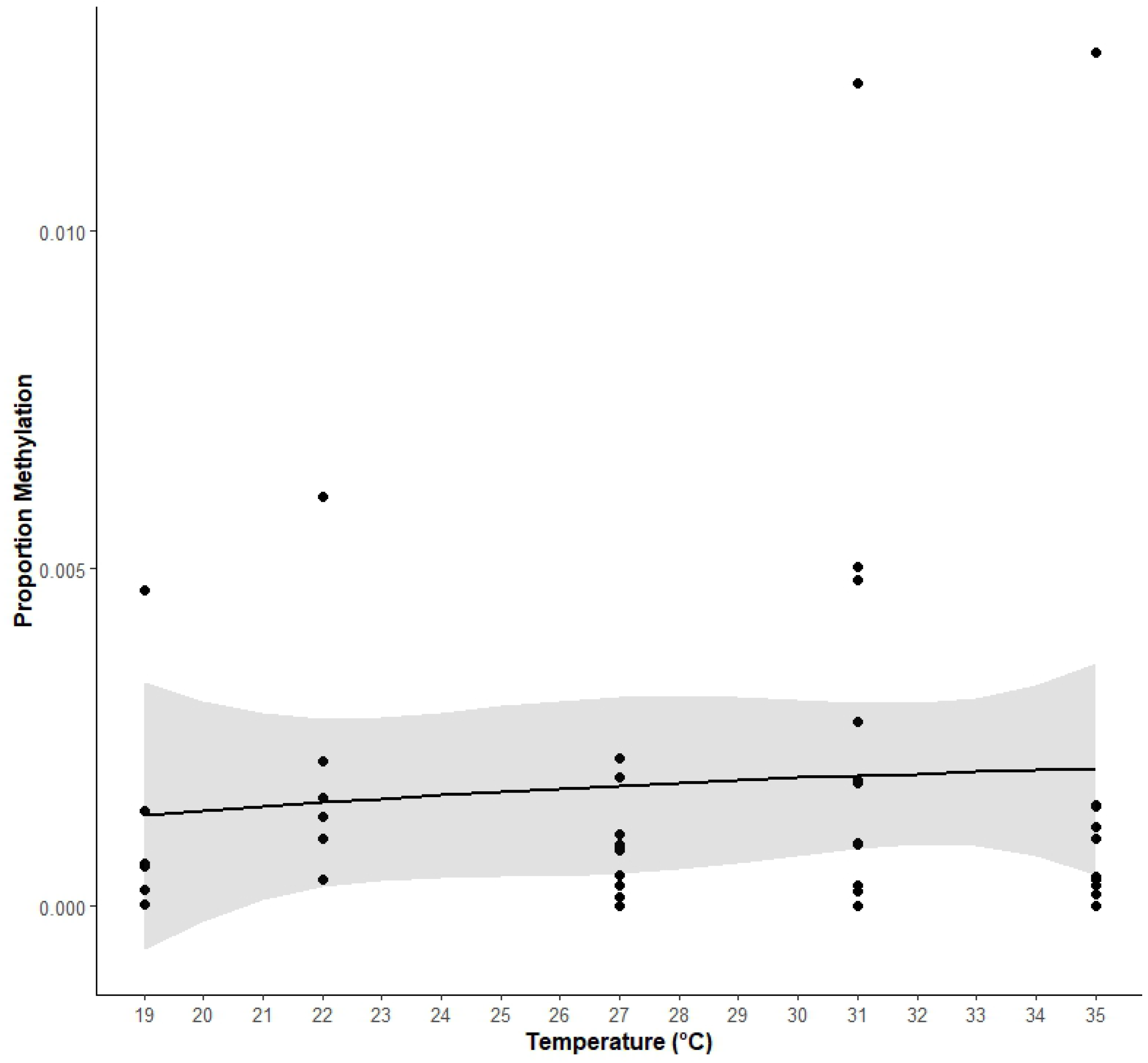
Proportion methylation levels in untreated female beetles across temperatures (*n* = 44). Data was plotted using a linear regression model with a quadratic coefficient for visualisation. Shaded regions represent 95% confidence intervals.

**S2 Table. Effects of treatment nested within temperature on methylation levels in a beta regression model controlling for block effects of ELISA plates (baseline = H2O, *n* = 64)**. For main effects of temperature in the model, see Results. Significant values are indicated in bold.

**S3 Table. Power to detect significant effects of 125nM 3AB or Zeb (nested in temperature, after controlling for plate effects) on whole-genome methylation at different simulated sample sizes**. Matrix entries represent our statistical power (ß) at α = 0.05. Values > 0.8 are in bold.

**S2 Fig.**
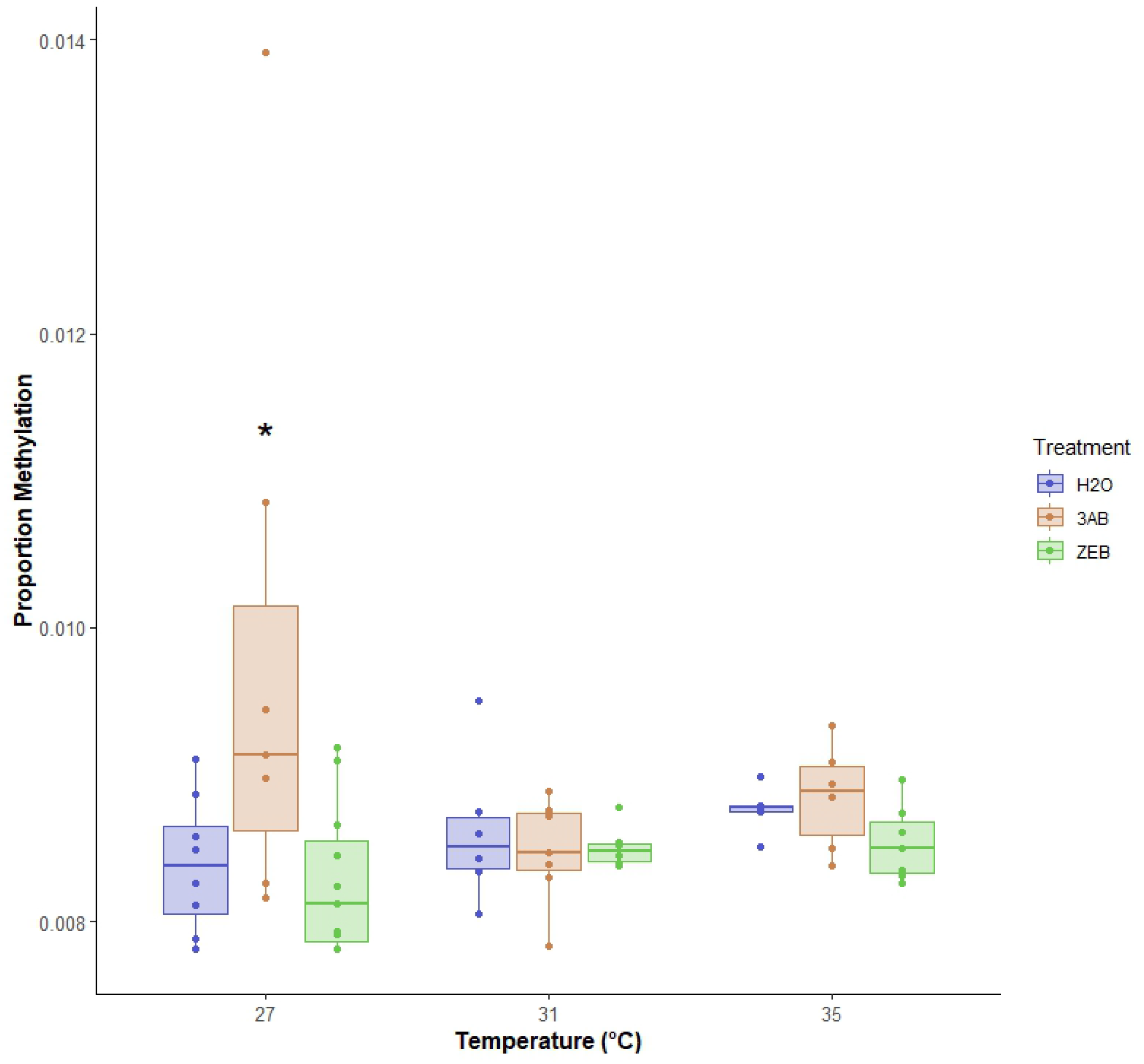
Proportion methylation levels in treated female beetles across 27°C, 31°C and 35°C (*n* = 64). Within each temperature: water = blue, left; 3AB = orange, middle; Zeb = green, right. Asterisks indicate significant values in comparison to water treatment within each temperature category. Each box represents the median, lower 25th and upper 75th percentile, with whiskers showing the values within 1.5 times the interquartile ranges. Outside values are >1.5 times and <3 times beyond the interquartile ranges.

**S3 Fig.**
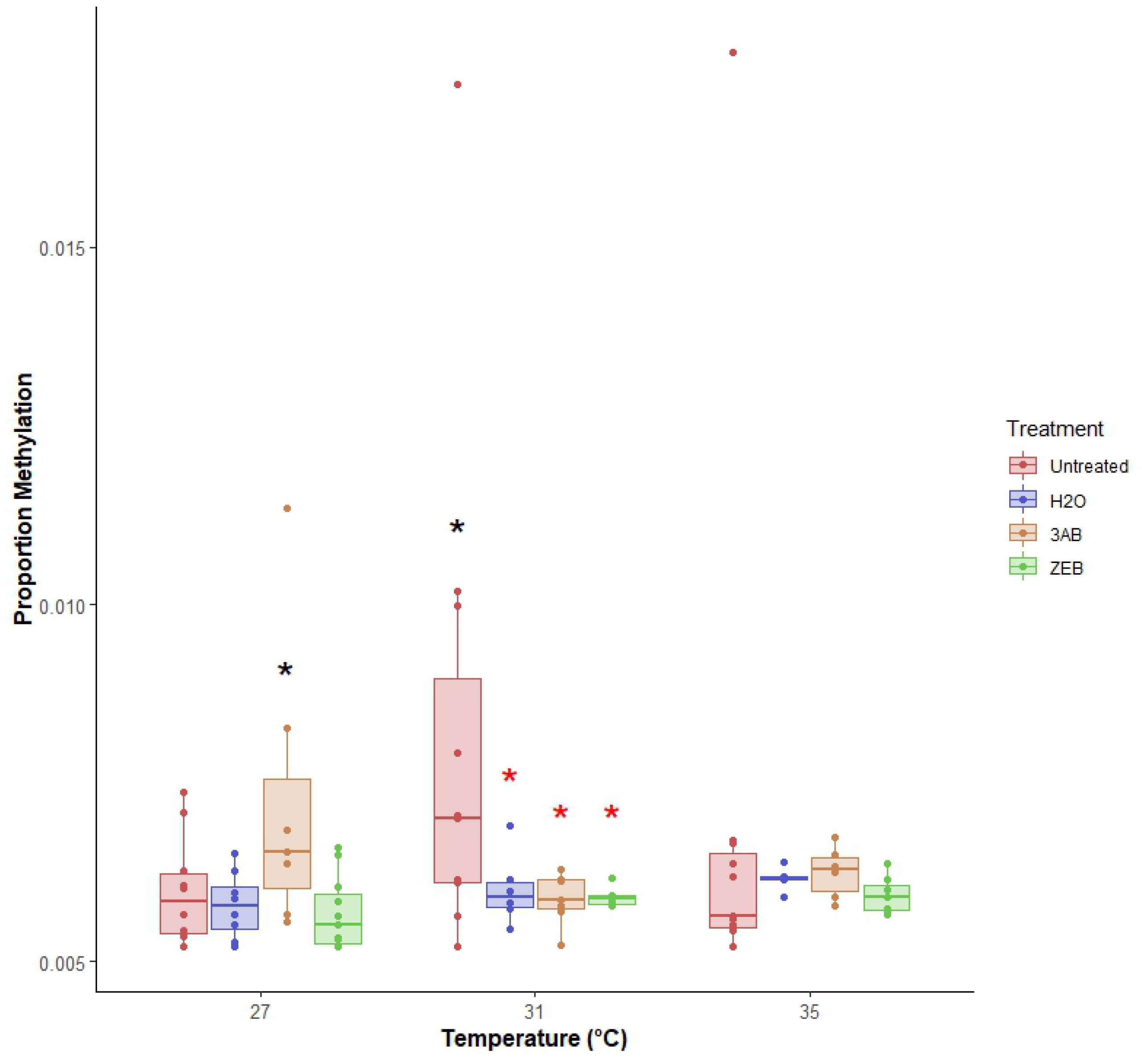
Proportion methylation levels in untreated and treated female beetles across 27°C, 31°C and 35°C (*n* = 96). Within each temperature: untreated = red, far-left; water = blue, middle-left; 3AB = orange, middle-right; Zeb = green, far-right. Black asterisks indicate significant values in comparison to water treatment within each temperature category. Red asterisks indicate significant values in comparison to untreated within each temperature category. Each box represents the median, lower 25th and upper 75th percentile, with whiskers showing the values within 1.5 times the interquartile ranges. Outside values are >1.5 times and <3 times beyond the interquartile ranges.

**S4 Table. Forward and reverse sequences of gene primers used for qPCR**. Custom primers for the transcripts of interest and reference transcripts were designed by Primerdesign Ltd and determined from the C. maculatus genome.

**S5 Table. Linear models explaining heat shock protein (*Hsp70* and *Hsp90*) and vitellogenin (*Vtg*) expression in whole adult males and females (baseline = Female, *n* = 6)**.

**S6 Table. Nested linear models explaining heat shock protein (*Hsp70* and *Hsp90*) and vitellogenin (*Vtg*) expression in adult male and female body parts (baseline = Female + Body, *n* = 11)**.

